# The impact of interspecies recombination on human herpes simplex virus evolution and host immune recognition

**DOI:** 10.1101/472639

**Authors:** Amanda M. Casto, Pavitra Roychoudhury, Hong Xie, Stacy Selke, Garrett A. Perchetti, Haley Wofford, Meei-Li Huang, Georges M. G. M. Verjans, Geoffrey S. Gottlieb, Anna Wald, Keith R. Jerome, David M. Koelle, Christine Johnston, Alexander L. Greninger

**Affiliations:** Department of Medicine, University of Washington, Seattle, Washington, USA; Department of Laboratory Medicine, University of Washington, Seattle, Washington, USA; Department of Viroscience, Erasmus MC, Rotterdam, The Netherlands; Research Center for Emerging Infectious and Zoonoses, University of Veterinary Medicine Hannover, Hannover, Germany; Department of Global Health, University of Washington, Seattle, Washington, USA; Center for Emerging and Re-Emerging Infectious Diseases, University of Washington, Seattle, Washington, USA; Vaccine and Infectious Diseases Division, Fred Hutchinson Cancer Research Center, Seattle, Washington, USA; Benaroya Research Institute, Seattle, Washington, USA

## Abstract

Among the most ubiquitous of human pathogens, HSV-1 and HSV-2 are distinct viral species that diverged about six million years ago. At least four ancient HSV-1 x HSV-2 interspecies recombination events have affected the HSV-2 genome, with recombinants and non-recombinants at each locus circulating today. Though interspecies recombination has occurred in the past, its importance in HSV evolution remains incompletely defined. Using 255 newly-sequenced and 219 existing HSV genome sequences, we comprehensively assessed interspecies recombination in HSV. The novel recombinants we identify demonstrate that the sizes and locations of interspecies recombination events in HSV-2 are more variable than previously appreciated. One novel recombinant arose in its current host, showing for the first time that interspecies recombination occurs in contemporary HSV populations. We also demonstrate that interspecies recombination affects T-cell recognition of HSV. Our findings indicate that interspecies recombination can significantly influence genetic variation in and host immunologic response to HSV-2.

## Introduction

The herpes simplex viruses (HSV-1 and HSV-2) are ubiquitous human pathogens with 3.7 billion HSV-1 and 417 million HSV-2 infected individuals worldwide (Looker et al., 2015a, 2015b). Both viruses establish lifelong infections typically characterized by mild, intermittent clinical symptoms. However, HSV can also cause significant morbidity and mortality, particularly among the immunocompromised and in neonates (Corey et al., 1983; Corey and Wald, 2009). Furthermore, genital HSV-2 infection has helped fuel the HIV epidemic by increasing the risk of HIV infection (Johnson et al., 2011; Masese et al., 2015; Zhu et al., 2009). While antivirals can reduce symptoms, they do not cure infection and do not completely prevent viral shedding and transmission, leading to the urgent need for an HSV vaccine (Gottlieb et al., 2016).

Instrumental in the development of an HSV vaccine and of new pharmaceutical therapies for HSV is a better understanding of the evolution of HSV and of the genetic variation among viral strains. HSV-1 and HSV-2 diverged from one another about 6 million years ago after which HSV-1 evolved in the human lineage and HSV-2 in the chimpanzee lineage (Wertheim et al., 2014). A human ancestor then acquired HSV-2 as a zoonotic infection from a chimpanzee ancestor 1.4 – 3 million years ago (Wertheim et al., 2014). Early studies of HSV genomes indicated that the viral species were relatively homogenous (mean pairwise distances among HSV-1 and HSV-2 strains are 0.8% and 0.2%, respectively) (Kolb et al., 2015). However, in 2015, a variant HSV-2 strain was described that was highly divergent from other HSV-2 samples at a single genomic locus (Burrel et al., 2015). This divergent region was likely affected by an ancient HSV-1 x HSV-2 interspecies recombination event with both recombinant and non-recombinant genotypes at this locus observed among HSV-2 strains today (Burrel et al., 2017; Koelle et al., 2017). Ultimately, four loci, totaling about 1% of the HSV-2 genome, were found that carried evidence of such recombination events (Burrel et al., 2017; Koelle et al., 2017). For three of the loci, subgenic regions within the UL29, UL30, and UL39 genes, recombinant genotypes were commonly observed among HSV-2 samples. Only one HSV-2 sample has been found that carries HSV-1 sequence at the fourth locus, which falls within UL15.

The high degree of divergence between recombinant and non-recombinant HSV-2 strains within these regions relative to the mean pairwise divergence elsewhere in the genome demonstrates that interspecies recombination could significantly contribute to variation among HSV-2 strains. However, numerous questions remain about the role of interspecies recombination in the evolution of HSV-2. In particular, it remains unknown whether interspecies recombination can affect loci other than those previously described in UL15, UL29, UL30, and UL39 and if interspecies recombination continues to affect contemporary HSV populations as all events described to date are thought to have occurred in a historical context. If HSV interspecies recombinants are still being generated, it is also unclear if the increasing number of immunocompromised hosts (Harpaz et al., 2016) or if changes in the epidemiology of HSV infection, such as the increase in genital HSV-1 (Chiam et al., 2010; Gilbert et al., 2011; Ryder et al., 2009), will alter the impact interspecies recombination has on genomic variation in HSV. We sought to answer these questions by performing a comprehensive survey of interspecies recombination on a large dataset of HSV genome sequences comprised of both previously available and newly-generated sequences.

## Results

### New sequencing more than doubles pool of available HSV genomes

We generated genome sequences for a total of 59 HSV-1 clinical samples, nearly doubling the number of HSV-1 genomes available for analysis (Supplementary Table 1). Fifty-eight of these samples were collected in Seattle, WA, USA, and one was from Uganda (Supplementary Table 2). Nine out of these newly-sequenced samples were from 6 persons who are genitally co-infected with HSV-1 and HSV-2 (Supplementary Table 3). For HSV-2, we generated 196 new genome sequences, increasing the total number of available HSV-2 genomes by 137% (Supplementary Table 4). Most (161 out of 196, or 82%) of these newly-sequenced samples were collected in Seattle, though samples collected in Cameroon (20), Peru (3), Senegal (10), and Uganda (2) were also sequenced. Eleven of these samples were from 6 persons genitally co-infected with HSV-1 and HSV-2. The countries of origin as well as the gender and HIV status of the carriers of viruses are reported for all newly-generated HSV genomes and for all the HSV genomes that were available in GenBank as of April 2018 in Supplementary Tables 1, 2, and 4.

### No evidence of interspecies recombination found within HSV-1 genomes

We examined the newly-sequenced HSV-1 genomes for HSV-1 x HSV-2 recombination events using both the Recombination Detection Program (RDP) software and manual alignment review (see Methods). All HSV-1 genomes were aligned with an HSV-2 reference (SD90e, KF781518) and a chimpanzee herpesvirus, or ChHV, reference (NC_023677) separately to avoid detection of intraspecies recombination events. As a positive control, we first reviewed the output of RDP for the regions of UL29 and UL30 where the HSV-2 reference strain SD90e carries recombinant HSV-1 sequence. RDP detected the UL29 and UL30 events in all 59 of the HSV-1-SD90e-ChHV alignments. RDP did return other putative events for all HSV-1 alignment trios. However, these events generated higher p-values than the UL29 and UL30 events for the same metric (Supplementary Note 1, Supplementary Figure 1) and we did not see evidence for recombination at the indicated loci on manual review. Overall, in line with previous results (Burrel et al., 2017; Koelle et al., 2017), we found no evidence of HSV-1 x HSV-2 recombination in the 59 newly-sequenced HSV-1 genomes.

### Interspecies recombination event spans multiple ORFs and is stable within host

Next, we analyzed 230 HSV-2 (196 newly generated and 34 previously unanalyzed) genome sequences for interspecies recombination again using both RDP and manual review. Five previously undescribed interspecies recombination events were observed. A sample collected from an HIV negative woman in Seattle (2015-14086, MF510363) contained 7 kilobases (kb) of HSV-1 sequence spanning half of UL29 and all of UL30 and UL31 (nucleotides 60,761 - 67,777 in SD90e reference) (Figure 1A, Supplementary Figure 2). The longest interspecies recombination event described prior to this measured 538 basepairs (bp) (Burrel et al., 2017; Koelle et al., 2017). Sanger sequencing confirmed the breakpoints of this event. We sequenced two additional samples from the same person (2013-34209, MH790566; 2014-13047, MH790604; Figure 1B). The 3 sequenced samples were collected over a period of 659 days. The 2 additional samples also contained the 7 kb UL29 – UL31 recombination event. Across the recombinant region, the 3 sequences were identical (only coding sequence was considered in sequence comparisons). The three HSV-1 sequences with the most similarity to the recombinant within the recombinant region were all collected in Seattle, including two that differed by 5 SNPs and one that differed by 6 (Figure 1C). Outside the recombinant region, there was only one polymorphic site among the three sequences; they formed a single clade in a phylogenetic tree of all available HSV-2 sequences (Figure 1D).

**Figure 1:**
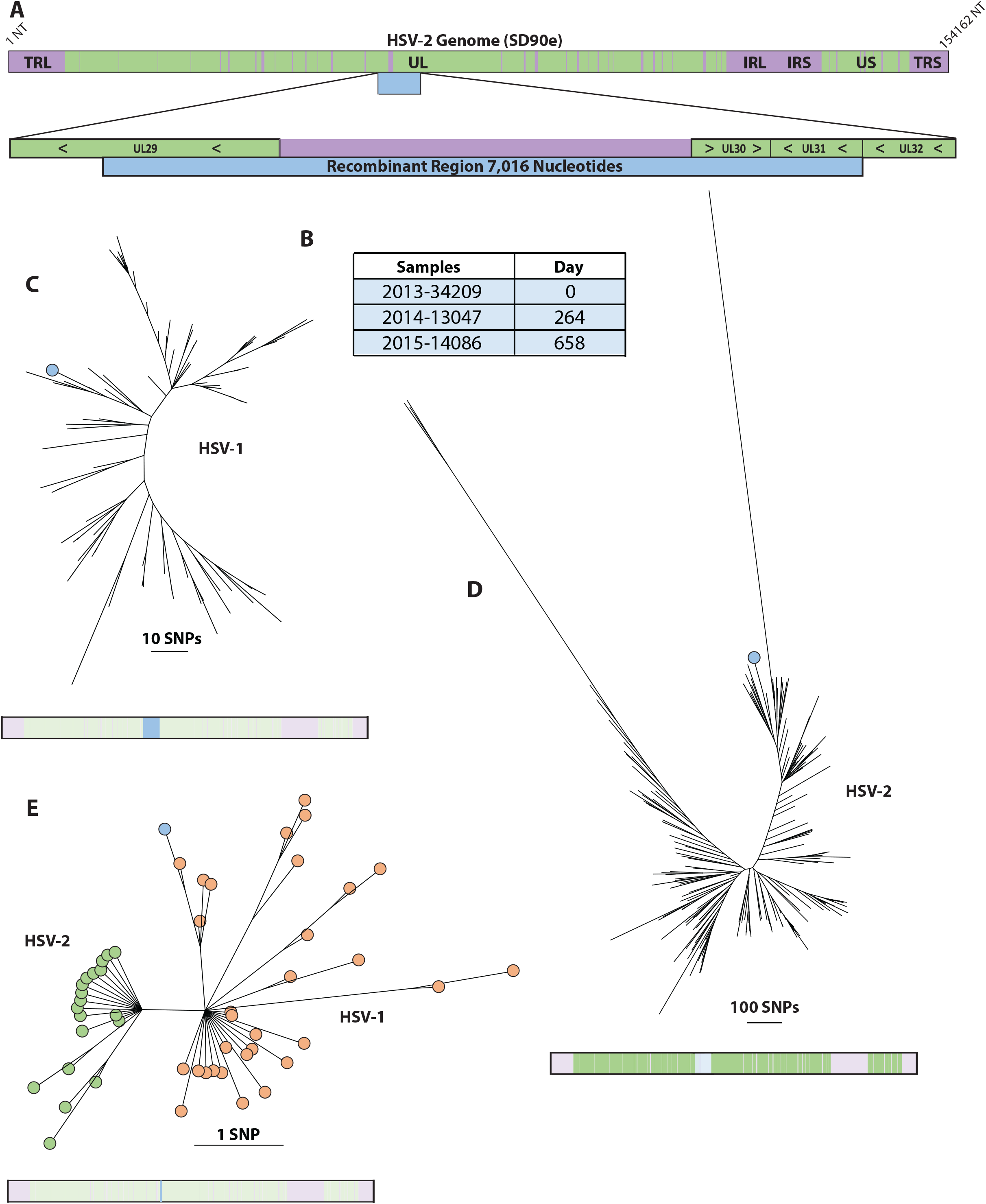
UL29 – UL31 Recombinant. A) Schematic showing the size and position of the recombination event relative to the HSV-2 genome and within its genic neighborhood. Green represents coding regions and purple represents non-coding regions. The recombinant region is represented by a blue bar. B) List of all sequenced samples collected from the person carrying the UL29 – UL31 recombinant with the day of sample collection relative to the collection date of the first sample. C) Phylogenetic tree of the recombinant region extracted from HSV-1 sequences and from the UL29 – UL31 recombinant. The bar below the tree shows the extracted region (colored in bright blue) relative to the rest of the genome (faded out). The branch representing the UL29-UL31 recombinants is marked with blue circle. D) Phylogenetic tree of all coding sequences (concatenated) excluding UL29, UL30, and UL31 extracted from HSV-2 sequences. The bar below the tree shows the extracted regions (colored in bright green) relative to the rest of the genome (faded out). All 3 HSV-2 sequences from the person carrying the UL29 – UL31 recombinant clustered on the branch marked with the blue circle. E) Phylogenetic tree of the UL30 recombinant region extracted from HSV-1 sequences, HSV-2 sequences with the UL30 CR genotype, and HSV-2 sequences with the UL29 – UL31 recombination event. The bar below the tree shows the extracted region (colored in bright blue) relative to the rest of the genome (faded out). The UL29 – UL31 recombinant branch is marked with a blue circle. HSV-1 branches are marked in orange and HSV-2 branches are marked in green.

Because it spans the entire UL30 ORF, this event encompasses the region affected by the previously-defined recombination event in UL30. It does not encompass the part of the UL29 ORF where the UL29 recombination event is observed (Burrel et al., 2017; Koelle et al., 2017). We hypothesized that the UL30 recombinant genotype could have descended from the UL29 – UL31 recombinant through backcrosses. However, in a phylogenetic tree of the UL30 recombinant region, the UL29 – UL31 recombinant does not cluster with samples carrying the UL30 recombination event, suggesting that the UL29 – UL31 recombinant and the UL30 recombinants were derived from different HSV-1 parental strains (Figure 1E).

### Interspecies recombinant generated in HSV-1/HSV-2 genitally co-infected host

A second novel interspecies recombination event was observed in a sample from a different HIV negative woman from Seattle (1996-26333, MH790638). This person has had both HSV-1 and HSV-2 detected in genital swabs. This event is more than 6 kb in length, spanning half of UL47 and all of UL48, UL49, and UL49A, and most of UL50 (nucleotides 102,443 to 108,479 in the SD90e reference) (Figure 2A, Supplementary Figure 3). Sanger sequencing also confirmed the breakpoints of this event. Within the recombinant region, the recombinant HSV-2 sample differed from an HSV-1 sample from the same person (1995-63175, MG999862) by just one SNP (Figure 2B). We then compared the recombinant region to the rest of our HSV-1 dataset. The next most similar sequence had 6 nucleotide differences and had been collected in Germany, followed by a sequence with 7 differences that was collected in Seattle. These data suggest that the HSV-1 strain collected from the person with the UL47-UL50 recombinant is most likely the HSV-1 parent of the recombinant strain (Figure 2C).

**Figure 2:**
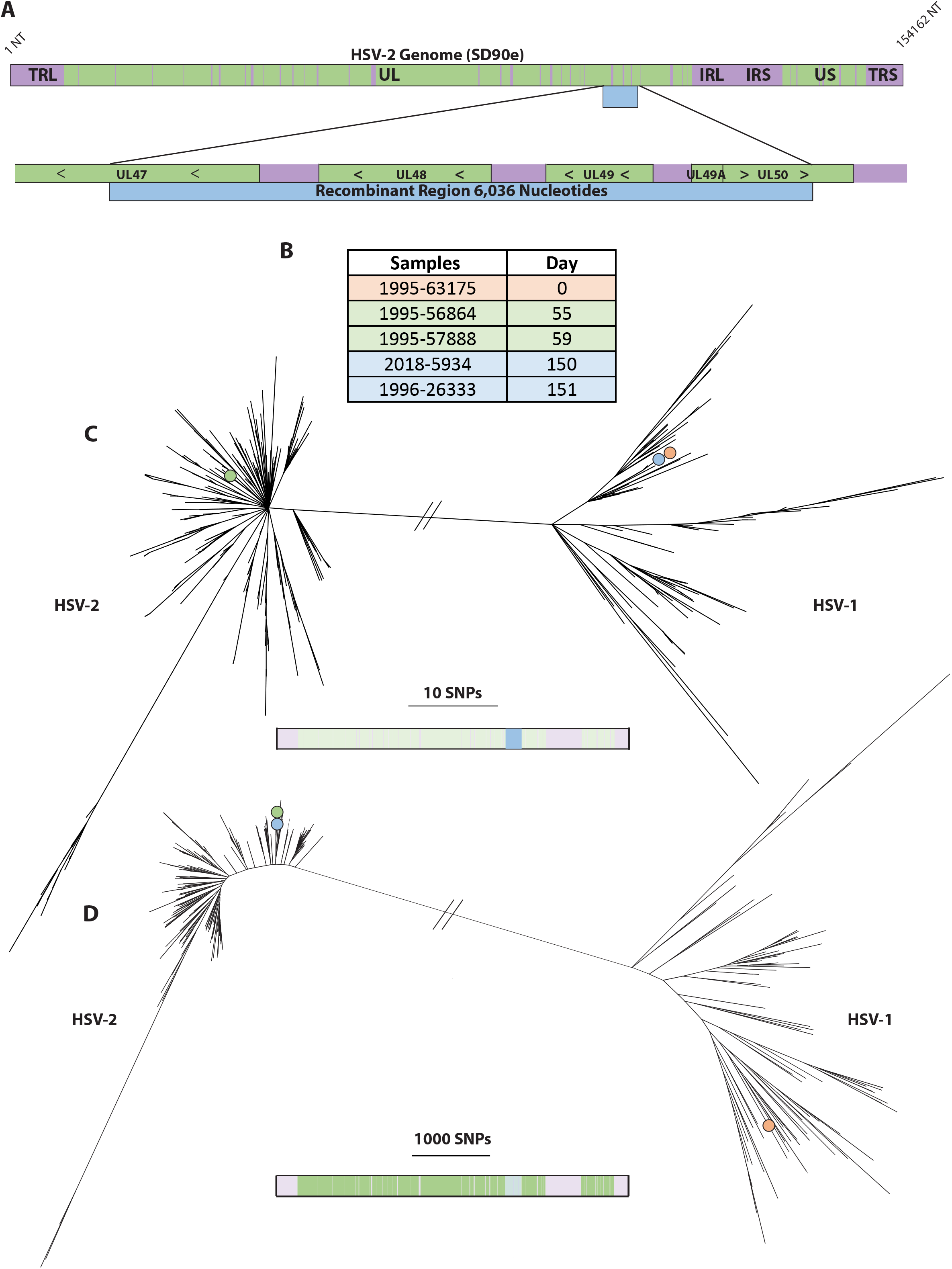
UL47 – UL50 Recombinant. A) Schematic showing the size and position of the recombination event relative to the HSV-2 genome and within its genic neighborhood. Green represents coding regions and purple represents non-coding regions. The recombinant region is represented by a blue bar. B) List of all sequenced samples collected from the person carrying the UL47 – UL50 recombinant with the day of sample collection relative to the collection date of the first sample. C) Phylogenetic tree of the recombinant region extracted from all HSV-1 and HSV-2 sequences. The bar below the tree shows the extracted region (colored in bright blue) relative to the rest of the genome (faded out). The branches marked with colored circles represent the position in the tree of the various samples collected from the person carrying the UL47 – UL50 recombinant. These colors correspond to those in the table in B. D) Phylogenetic tree of all coding sequences (concatenated) excluding UL47, UL48, UL49, UL49a, and UL50 extracted from all HSV sequences. The bar below the tree shows the extracted regions (colored in bright green) relative to the rest of the genome (faded out). The branches marked with colored circles represent the position in the tree of the various samples collected from the person carrying the UL47 – UL50 recombinant. These colors correspond to those in the table in B.

We next sequenced three additional HSV-2 samples collected before the original sample (1996-26333) from the same person (1995-56864, MH790588; 1995-57888, MH790664; 2018-5934, MH790582). The samples (4 HSV-2 and one HSV-1) sequenced for this person were collected over a 151-day period (see Figure 2B). One of the 3 additional HSV-2 samples (2018-5934), collected one day before the original sample (1996-26333), carried the UL47-UL50 recombination event. However, two other samples (1995-56864, 1995-57888) both collected earlier than the recombinant HSV-2 samples did not contain the recombination event. Outside the recombinant region all four of the HSV-2 samples phylogenetically clustered together (Figure 2D), suggesting that that the non-recombinant HSV-2 strain from this person is the most likely HSV-2 parent for the UL47 – UL50 recombinant.

### Novel events vary widely in size and location

In addition to the two large recombination events, we observed three other small novel events. The first of these was a 40 bp event in UL17 (Supplementary Figure 4), which resulted in 9 nucleotide and 3 amino acid changes. This was observed in a single sample from a man with HIV infection from Seattle (2005-42278, MH790643). The second small event was a 52 bp event in UL28 that resulted in 4 nucleotide and no amino acid changes. It was observed in 4 samples from 4 different individuals, all of which were collected in East Africa (three in Uganda, one in Kenya) (2012-18385, MF510324; 2009-3495, MF510280; 2012-18420, MF510337; 2009-4463, MF621258). This small event was detected only by manual review. Finally, we observed a 259 bp event in UL32 which resulted in 25 nucleotide and 4 amino acid changes (Supplementary Figure 5). This event was observed in 3 samples collected over a 12-year period from the same HIV negative man in Seattle (2002-14972, MH790585; 2014-14807, MF510306; 2014-14811, MF510292).

### Four rare genotypes found at UL29 recombinant locus

We next sought to characterize variation at the 3 loci in HSV-2 (UL29, UL30, and UL39) where interspecies recombinant genotypes are common (Burrel et al., 2017; Koelle et al., 2017) by assigning a genotype to each HSV-2 sequence for each of these 3 loci (Supplementary Note 2). We report first on the results for UL29, which encodes the HSV single stranded binding protein, ICP8 (Mapelli et al., 2005) (Figure 3A). The interspecies recombination event in UL29 affects a 121 amino acid (363 bp) region in the C-terminal half of the protein. The full length recombinant genotype, which we called the common recombinant or CR genotype, was present in 98.1% of HSV-2 sequences (209 out of 213) (see methods for a description of the sequences included in this count). Only one sequence out of 213 (0.5%) carried the non-recombinant (NR) genotype (Figure 3B). This HSV-2 sample was collected from an HIV positive woman in Cameroon. We additionally noted a total of 4 rare recombinant genotypes at this locus, 3 of which are previously undescribed (the fourth rare genotype is described in Koelle et al., 2017). Each of these rare genotypes was seen in a single sample from Cameroon (R1), Seattle (R2), DRC (R3), and Kenya (R4). The CR genotype at UL29 is characterized by two stretches of HSV-1 sequence separated by a non-recombinant region and as such has 4 different breakpoints (UL29 nucleotide positions 2,076 and 2,136 mark of the ends of the HSV-1 sequence for the first block; 2,196 and 2,439 mark the ends of the second block). All of the rare genotypes share at least 2 breakpoints with the CR genotype except for R1, which has just one stretch of HSV-1 sequence with 2 unique breakpoints (UL29 nucleotides 1,815 and 2,196). Intriguingly, the recombinant portion of UL29 maps to functionally critical portions of the protein which allow it to resist proteolysis and interact with single-stranded DNA (Mapelli et al., 2005) (Figure 3C).

**Figure 3:**
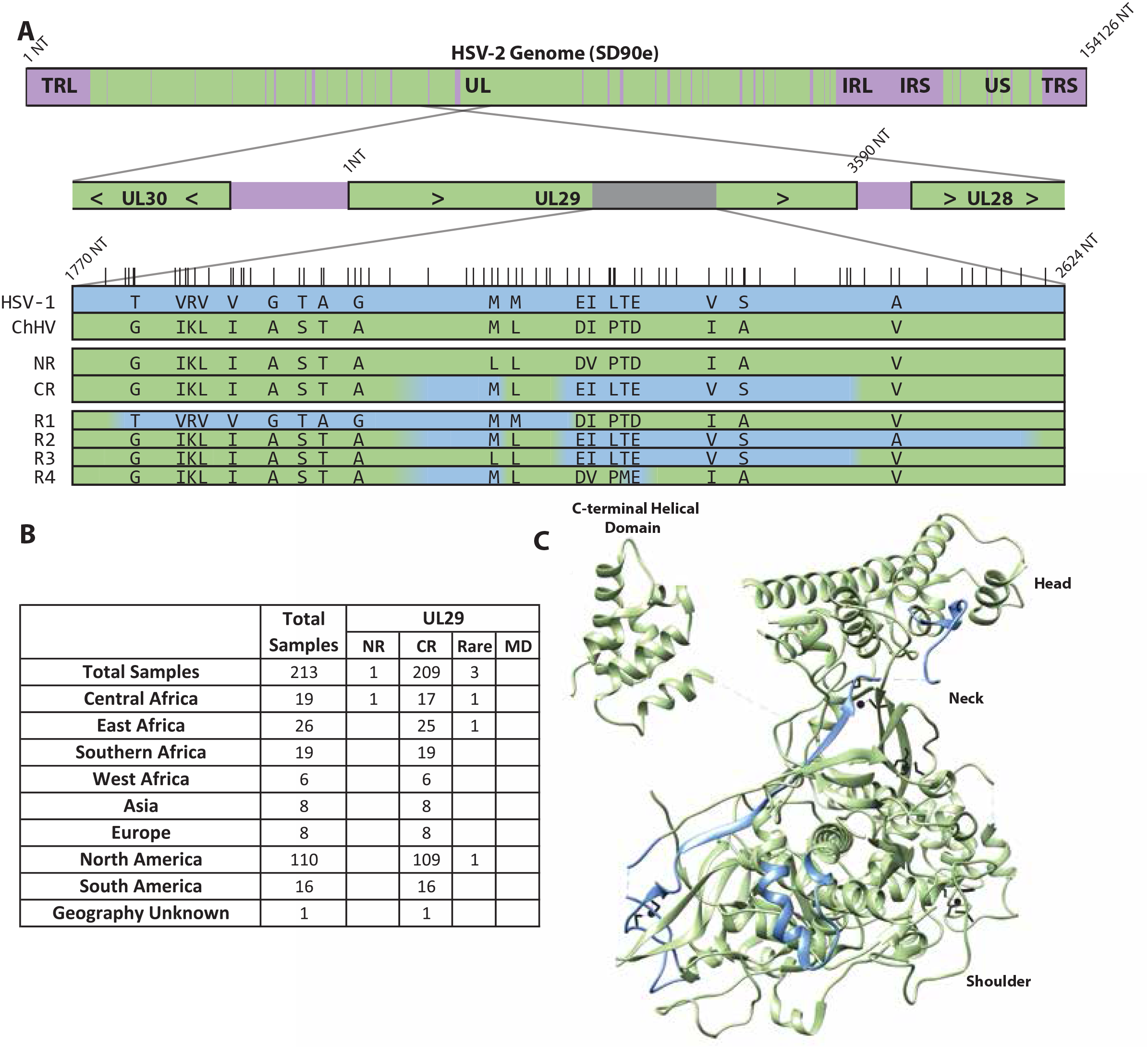
UL29 Recombinant Region. A) Schematic showing the HSV-1 sequence blocks for each UL29 recombinant genotype. HSV-1 sequence is colored in blue. ChHV/non-recombinant HSV-2 coding sequence is colored in green. Intergenic regions are purple. Letters denote amino acids that differ among the various genotypes and vertical lines above the HSV-1 row denote the location of nucleotides that differ among the various genotypes (“genotype-defining” SNPs, see Supplementary Note 2). The top bar represents the entire HSV-2 genome while the second bar represents the region of the genome containing UL29. Of note, the UL29 gene is oriented in the reverse direction. The orientation of the second bar and the genotype bars are flipped relative to the full genome bar. B) Number of samples from each geographic region with the NR, CR, and rare genotypes. MD denotes samples with missing data. Note that samples LT797622 – LT797636, LT797682, LT797786, LT799380 (samples from Burrel et al., 2017) were not included in these counts. C) Crystal protein structure of the UL29 gene product, ICP8, the HSV single stranded DNA binding protein. The part of the protein encoded by the recombinant region is highlighted in blue.

### UL30 has two recombinant genotypes

UL30 encodes the catalytic subunit of the DNA polymerase (Liu et al., 2006). The interspecies recombination event in UL30 affects a 180 amino acid (538 bp) region in the C-terminal half of the protein (Figure 4A). The full length or common recombinant (CR) genotype was observed in 95.7% (202 out of 211 with 2 excluded for missing data) of HSV-2 sequences (Figure 4B). The non-recombinant genotype (NR) was observed in 8 sequences (3.8%). These sequences were from Uganda (4), Cameroon (2), and Seattle, WA, USA (2). Two (10.5%) out of 19 sequences from Central Africa and four (15.3%) out of 26 sequences from East Africa had the NR genotype. There was only one rare genotype (R1) observed for UL30. This genotype has not been described previously and has a shorter HSV-1 block than the common recombinant, with a unique 5’ breakpoint (UL30 nucleotide 2,926) but the same 3’ breakpoint (UL30 nucleotide 3,409) as the CR genotype. The R1 genotype has no amino acid differences relative to the CR genotype. The sample carrying R1 at UL30 was collected in Seattle. Five other samples from the same person also carried this rare genotype.

**Figure 4:**
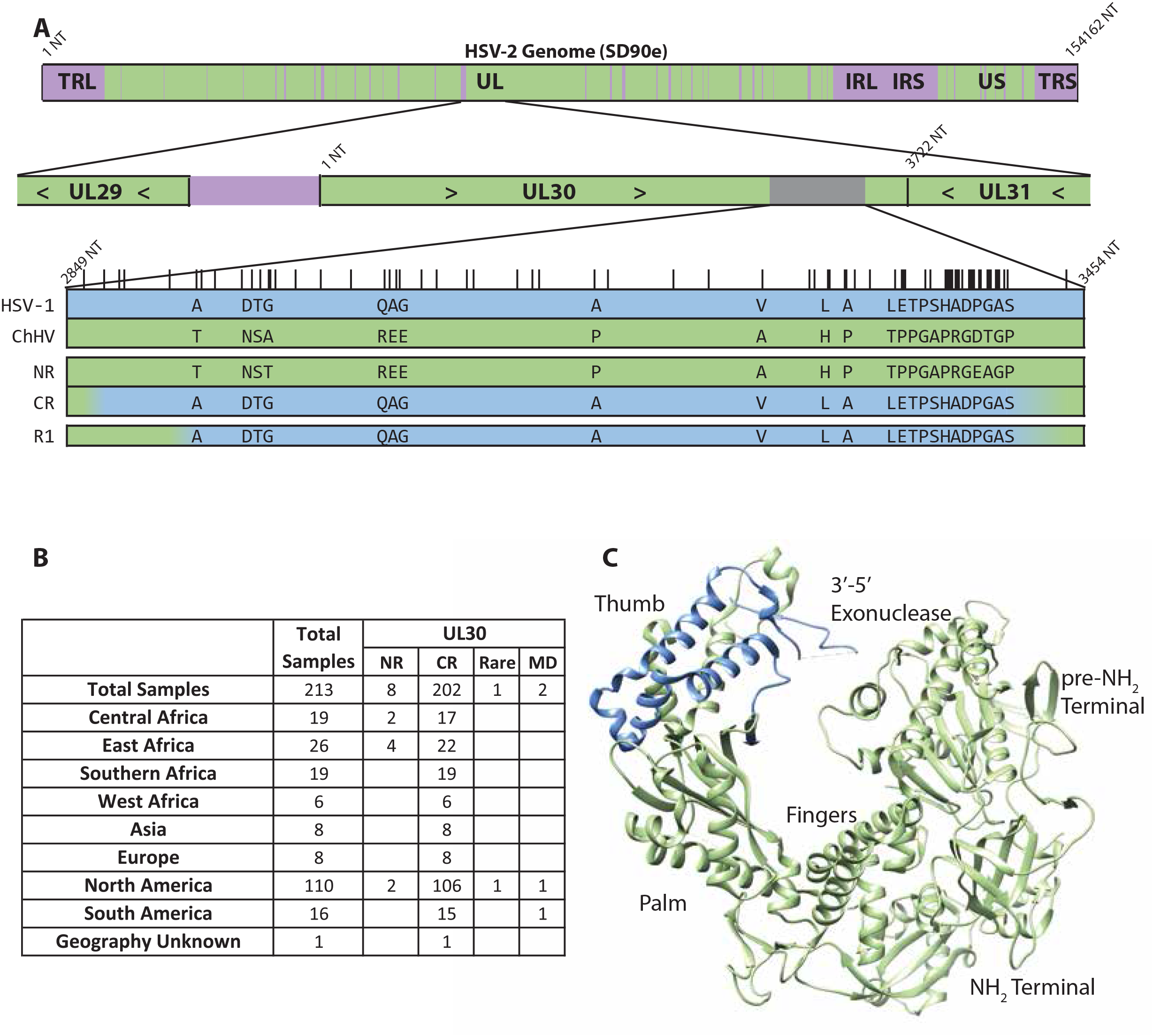
UL30 Recombinant Region. A) Schematic showing the HSV-1 sequence blocks for each UL30 recombinant genotype. HSV-1 sequence is colored in blue. ChHV/non-recombinant HSV-2 coding sequence is colored in green. Intergenic regions are purple. Letters denote amino acids that differ among the various genotypes and vertical lines above the HSV-1 row denote the location of nucleotides that differ among the various genotypes. The top bar represents the entire HSV-2 genome while the second bar represents the region of the genome containing UL30. B) Number of samples from each geographic region with the NR, CR, and rare genotypes. MD denotes samples with missing data. Note that samples LT797622 – LT797636, LT797682, LT797786, LT799380 (samples from Burrel et al., 2017) were not included in these counts. C) Crystal protein structure of the UL30 gene product, the HSV DNA polymerase. The part of the protein encoded by the recombinant region is highlighted in blue.

The recombinant region of UL30 encodes a portion of the thumb domain of the DNA polymerase (Liu et al., 2006) (Figure 4C). This critical domain interacts with double-stranded DNA as it leaves the catalytic center of the enzyme. Though none of the 23 residues affected by the recombination event have been associated with drug resistance to date, 21.5% of drug-resistance (acyclovir, penciclovir, foscarnet, cidofovir) mutations identified in the HSV-1 DNA polymerase are in the thumb domain (Sauerbrei et al., 2016; Topalis et al., 2016).

### UL39 contains a complex recombination locus

UL39 encodes the large subunit of the HSV ribonucleotide reductase (Conner et al., 1994). The interspecies recombination event affects a 152 amino acid (456 bp) region of this gene. This recombinant locus exhibits high sequence diversity with several common and numerous rare genotypes (Koelle et al., 2017). We defined genotypes at this locus first at the nucleotide level (see Supplementary Note 2). However, given the large number of nucleotide genotypes observed (21), we ultimately elected to group genotypes together if they had the same amino acid sequence. The most common nucleotide sequence corresponding to each amino acid sequence is illustrated in Figure 5A.

**Figure 5:**
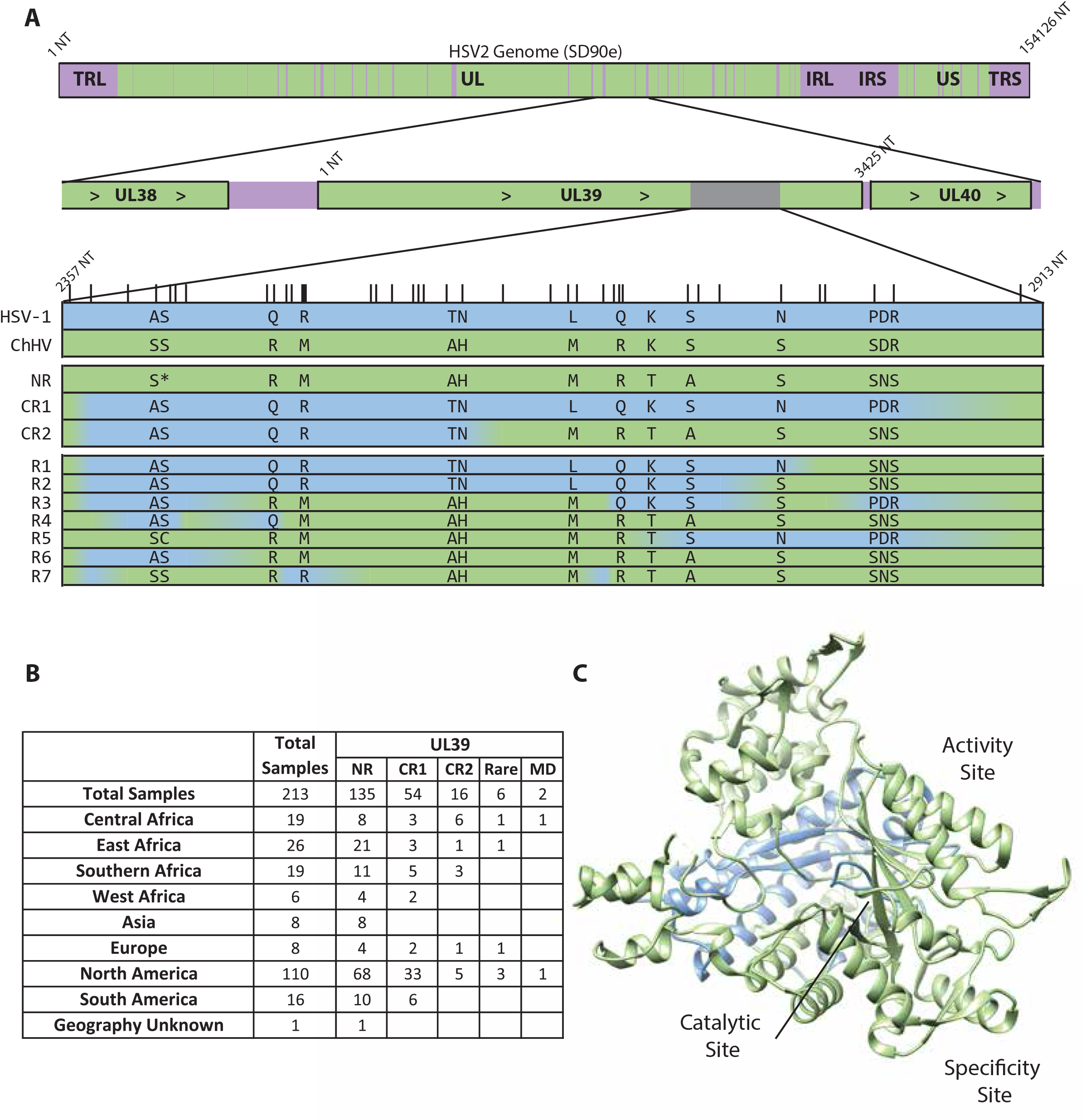
UL39 Recombinant Region. A) Schematic showing the HSV-1 sequence blocks for each UL39 recombinant genotype. HSV-1 sequence is colored in blue. ChHV/non-recombinant HSV-2 coding sequence is colored in green. Intergenic regions are purple. Letters denote amino acids that differ among the various genotypes and vertical lines above the HSV-1 row denote the location of nucleotides that differ among the various genotypes. The top bar represents the entire HSV-2 genome while the second bar represents the region of the genome containing UL39. Asterix (*) in NR genotype can be amino acid S or C depending on the sample. B) Number of sequences from each geographic region with the NR, CR, and rare genotypes. MD denotes sequences with missing data. Note that samples LT797622 – LT797636, LT797682, LT797786, LT799380 (sample from Burrel et al., 2017) were not included in these counts. C) Crystal protein structure of the UL39 gene product, the ribonuclease reductase (structure is of yeast ribonucleotide reductase). The part of the protein encoded by the recombinant region is highlighted in blue.

Most HSV-2 samples (135 out of 211, or 64.0%) had the non-recombinant (NR) genotype at UL39 (Figure 5B). There are 2 common recombinant genotypes at the UL39 locus. Common recombinant 1 (CR1) has 456 bp of HSV-1 sequence (breakpoints at UL39 nucleotides 2,373 and 2,829) and was seen in 54 (25.6%) of 211 (two with missing data excluded) samples. Common recombinant 2 (CR2) has 234 bp of HSV-1 sequence (breakpoints at UL39 nucleotides 2,373 and 2,607) and was seen in 16 (7.6%) of 211 samples. The remaining seven samples have 7 distinct, rare recombinant genotypes, 5 of which are previously undescribed (R3 and R7 described in Koelle et al., 2017). Samples with rare genotypes at UL39 were from the US (3), Cameroon (1), the DRC (1), Finland (1), and Kenya (1). Interestingly, the samples from Cameroon and from Kenya both also had rare genotypes for the UL29 locus.

Given the lack of a crystal structure for the HSV ribonuclease reductase, the protein structure of yeast ribonucleotide reductase (see Methods) is shown in Figure 5C (Xu et al., 2006). Three of the main murine subdominant epitopes recognized by HSV-1 specific CD8 cells have been mapped to the UL39 gene product (St Leger et al., 2011). One of these epitopes, which includes HSV-1 UL39 amino acids 822 – 829 (corresponding to HSV-2 UL39 amino acids 826 – 833), is within the region affected by interspecies recombination (St Leger et al., 2011). The NR version of this epitope contains two amino acid changes relative to the CR1 and CR2 genotypes. Functional experiments have demonstrated that the HSV-1 specific CD8+ T-cells that recognize the CR1/CR2 version of this epitope do not recognize the NR version (Salvucci et al., 1995).

### Recombinant genotypes at UL29, UL30, and UL39 are stable over time

To determine whether genotypes at the UL29, UL30, and UL39 loci are stable over time within hosts, we identified people who had more than one sequenced HSV-2 sample. Out of 65 such persons, the samples for 57 (87.7%) had the same genotypes at the UL29, UL30, and UL39 loci. The remaining 8 persons had all previously been identified as being infected with at least 2 distinct HSV-2 strains (Johnston et al., 2017).

One of these 8 persons, a man with HIV infection from Seattle, had a total of 4 sequenced samples. The two sequences that were analyzed in the Johnston et al., 2017 study (2003-18061, MF510298; 2007-22031, MF510341) had the same genotypes at the UL29, UL30, and UL39 loci with the NR genotype at the UL39 locus. However, we noted that a third sequence (2003-16029, KX574903) from this person had the CR1 genotype at UL39. We confirmed UL39 genotype assignments for all three of these samples using Sanger sequencing. We then compared these 3 sequences across the entire length of the genome. The two previously studied sequences differed from one another at 251 SNPs while the third sequence differed from the first two at 105 and 120 SNPs, respectively. In the Johnston et al. study, the strains in persons infected with two strains differed from one another by 87 to 305 SNPs (range based on 13 sequence pairs). Taken together, this suggests that this person is infected with at least 3 different HSV-2 strains.

### One HSV-2 sequence from Cameroon had no observed HSV-1 recombination events

We noted that one sample from Cameroon had the non-recombinant genotype at the UL29, UL30, and UL39 loci and did not contain any other interspecies recombination events. This sequence (2006-16150, MH790600) would be a useful alternative reference sequence to SD90e (KF781518) and HG52 (JX112656) for studies of HSV evolution as both of these sequences contain recombinant HSV-1 sequence. This sample has been sequenced to high quality with 50x or higher coverage for 95.8% of the full genome.

### Interspecies recombination impacts recognition of HSV by T-cells

In order to understand the functional importance of HSV interspecies recombination *in vivo*, we investigated whether interspecies recombination alters HSV recognition by CD4 and CD8 T-cells. We found that a polyclonal CD4 T-cell line (from an HSV-1-infected donor, GU14669) (Supplementary Table 5) strongly recognized full-length UL30 from both an HSV-1 strain (lab strain E115) and HSV-2 strains with the UL30 CR genotype (strains HG52 and 186, JX112656 and JN561323, respectively). However, these T-cells failed to recognize the NR version of UL30 (2008_15116, MF621257) (Figure 6A). We next examined T-cell recognition of the UL47-UL50 recombinant (1996_26333, MH790638) relative to HSV-1 and HSV-2 samples without the recombination event. A CD8 T-cell clone (1874.1191.22) (Supplementary Table 5) recognizing an epitope in HSV-2 UL47 laying outside the UL47 – UL50 recombination event (amino acids 551-559) but not recognizing HSV-1 recognized the UL47 – UL50 recombinant (Figure 6B). However, a CD8 T-cell clone (5101.1999.23) recognizing an epitope in HSV-2 UL47 at amino acids 289 – 298, which lies within the HSV-1 portion of UL47 in the recombinant, failed to recognize the recombinant. Conversely, two CD8 cell lines (TG3 UL48 N, TG3 UL48 C) that recognize HSV-1 UL48 but do not recognize HSV-2 UL48 gained novel recognition of the HSV-2 UL47-UL50 recombinant (Figure 6C, 6D, 6E). As a negative control, we also tested a CD4 cell clone (9447.28) that recognizes a peptide in UL49 identical in HSV-1 and HSV-2. As expected, this clone recognized the recombinant (Figure 6F). These results illustrate that interspecies recombination can serve as a mechanism of T-cell immune evasion.

**Figure 6:**
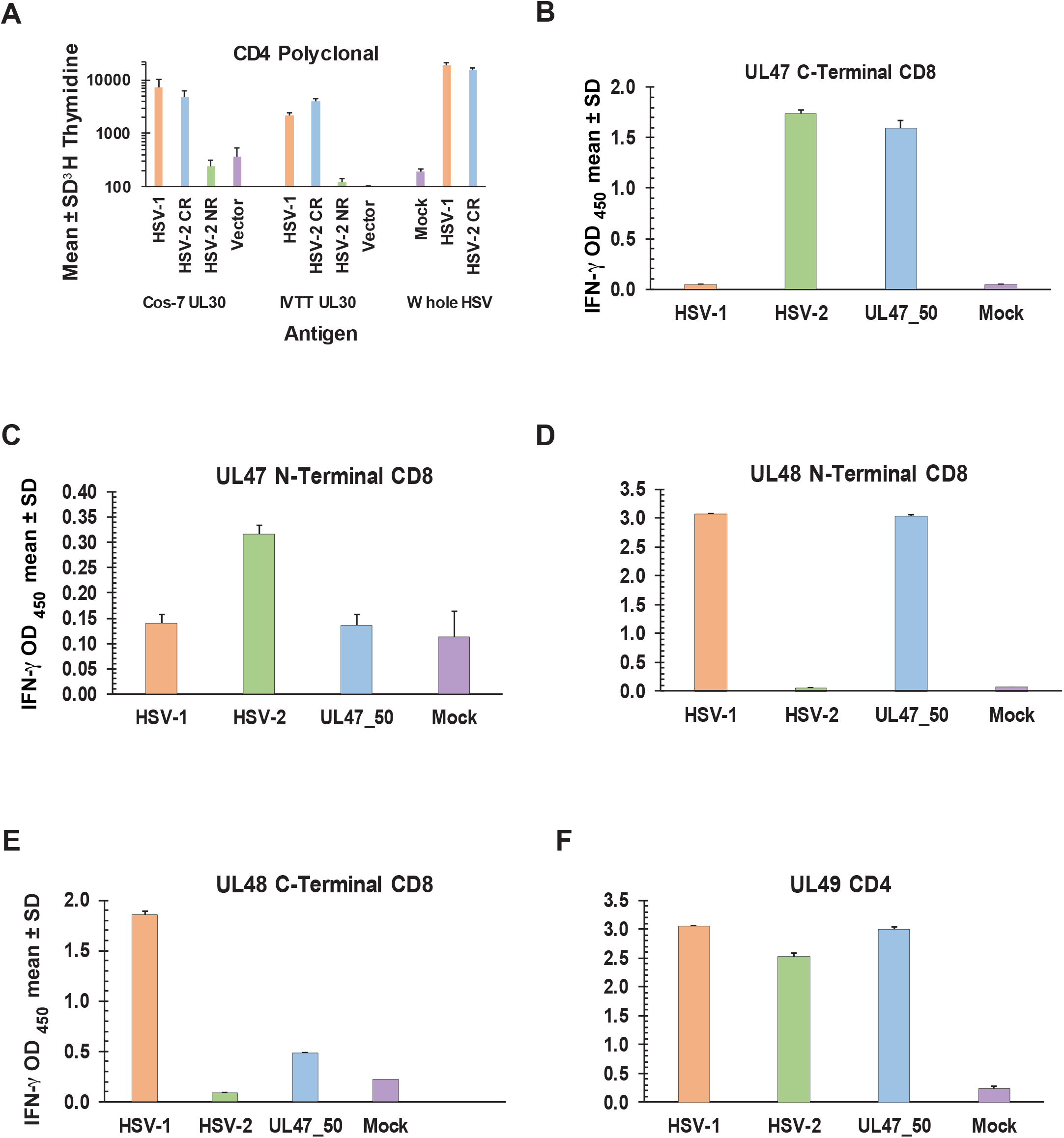
Differential recognition of HSV-1 x HSV-2 interspecies recombinants by HSV-specific T-cells. A) Polyclonal HSV-1-reactive CD4 T-cells from HSV-1-infected persons recognize UL30 expressed in either Cos-7 or IVTT systems from HSV-1 and HSV-2 UL30 CR, but not from HSV-2 UL30 NR genotype. Readout is T-cell proliferation. B) UL47 C-Terminal epitope-specific CD8 T-cell clone recognizes HSV-2 UL47-UL50 recombinant and non-recombinant HSV-2 but not HSV-1. C) UL47 N-Terminal epitope-specific CD8 T-cell clone recognizes non-recombinant HSV-2 but not HSV-1 or HSV-2 UL47-UL50 recombinant. D) UL48 N-terminal epitope-specific CD8 T-cell line recognizes HSV-1 and HSV-2 UL47-UL50 recombinant but not non-recombinant HSV-2. E) UL48 C-terminal epitope-specific CD8 T-cell line clone recognizes HSV-1 and HSV-2 UL47-UL50 recombinant but not non-recombinant HSV-2. F) UL49-specific CD4 T-cell clone recognizes HSV-1, non-recombinant HSV-2, and the HSV-2 UL47-UL50 recombinant. Data are secreted IFN-γ. All assays are triplicate.

## Discussion

It has long been known that HSV-1 and HSV-2 could recombine *in vitro* (Halliburton, 1980; Morse et al., 1977), but only recently appreciated that interspecies recombination has impacted the genome of nearly all HSV-2 clinical samples sequenced to date. Because HSV-2 has the potential to evolve more rapidly via recombination than through point mutation, a greater understanding of when, where, and why *in vivo* HSV interspecies recombination occurs and what its impacts are on human immunological response to HSV has important clinical and public health implications. Here we show that interspecies recombination events are more pervasive in the HSV-2 genome than previous appreciated and that they continue to be generated in circulating HSV-2 strains. We also demonstrate that such recombination can alter the recognition of HSV by CD4 and CD8 T-cells.

Consistent with earlier findings, we found no evidence of HSV-2 sequence in any of the HSV-1 genomes that we analyzed. While interspecies recombination events in HSV-1 may still exist, a profound asymmetry between the receptivity of the HSV-1 and HSV-2 genomes to sequence from the other HSV species is evident, given that almost all HSV-2 genomes contain some HSV-1 sequence. Since HSV-1 interspecies recombinants have been observed *in vitro*, the barrier to the presence of HSV-2 DNA in HSV-1 genomes is presumably downstream from the recombination event itself. It has been purposed that this barrier is at the level of transmission (Burrel et al., 2017). However, the failure to detect even one instance of recombinant HSV-1 in 135 HSV-1 genomes, including 9 from genitally co-infected persons (where the viruses are not relegated to different anatomic sites) is suggestive that the barrier may, in fact, be earlier in the HSV-1 life cycle. Overall, the one-way exchange of DNA between HSV-1 and HSV-2 *in vivo* is a curious feature of the evolution of these two viruses which does not yet have a clear explanation.

In HSV-2, recombinant genotypes at the UL29, UL30, and UL39 are observed at high frequencies, suggesting they have a selective advantage. Given the scarcity of the UL29 and UL30 NR genotypes, the recombinant regions in these two essential genes may prove useful as drug or vaccine targets given the sequence identity between HSV-1 and most HSV-2 strains at these loci. Our analysis of the UL29, UL30, and UL39 recombinant regions also established that there are both common and rare recombinant genotypes at all three loci. This finding suggests that these recombinant genotypes are not static but continue to evolve both through inter- and intraspecies recombination. Indeed, this process may be accelerating as rare genotypes, such as the NR genotypes at UL29 and UL30, are shuffled around the world by globalization. The potential for rapid creation of new genotypes through recombination at these loci is perhaps most concerning at the UL30 locus, as the region of the gene were HSV-1 sequence is commonly observed is also where about 20% of all described drug resistance mutations in UL30 are located (Sauerbrei et al., 2016; Topalis et al., 2016).

We also present significant additional data that the effects of interspecies recombination on HSV-2 are not restricted to the previously defined events in UL15, UL29, UL30, and UL39. Among the novel events we describe are two that are several kilobases in length and span multiple genes; both of these events are more than 10 times larger than the largest event described in Burrel et al., 2017 and Koelle et al., 2017. Furthermore, we provide evidence that these recombinant viruses are stable and can be persistently shed for years. Even if recombinant HSV-2 with such large recombination events are rare, any advantageous HSV-1 alleles in the region could theoretically disperse throughout the HSV-2 population via subsequent intraspecies recombination.

Our findings also challenge the supposition that new interspecies recombinant genotypes are no longer being generated in modern HSV-2 populations. In particular, the large sizes of the UL29-UL31 and UL47-UL50 events are by themselves suggestive that they were generated recently as intraspecies recombination has not had time to reduce their size through backcrosses with non-recombinant genomes. Furthermore, we present evidence that one of these two events, UL47-UL50, arose in the person from whom it was collected. This particular person is one of six genitally co-infected people from whom we have a sequenced HSV-2 sample. The high incidence of recombinant HSV-2 with large HSV-1 blocks in genitally co-infected persons is striking relative to the incidence of such recombinants among all individuals with a sequenced HSV-2 sample (1 in 6 compared to 2 in 231). This finding, coupled with our documentation of the generation of the UL47 – UL50 recombinant in a genitally co-infected person, suggests that genital co-infection fosters interspecies recombination. This is noteworthy as the incidence of genital HSV-1 has been shown in multiple studies to be increasing (Chiam et al., 2010; Gilbert et al., 2011; Ryder et al., 2009). The evidence we present here indicates that this epidemiologic shift could lead to an increase in the frequency of recombinant HSV-2.

The existence of interspecies HSV-2 recombinants has numerous potential impacts on clinical phenotype and on the development of new therapeutics, including an HSV vaccine. We demonstrate here that interspecies recombination can profoundly alter T-cell recognition of HSV. In particular, we showed that HSV-2 strains can completely gain or lose recognition by a particular T-cell clone through a single interspecies recombination event suggesting a mechanism by which such recombination could result in immune escape. Given the ability of interspecies recombination to significantly alter the T-cell repertoire that recognizes an HSV-2 strain, a better understanding of the natural history of HSV recombination and of the worldwide distribution of interspecies recombinants is absolutely crucial to the development of an effective HSV-2 vaccine. Other curative therapeutic approaches for HSV such as the disruption of latent HSV by targeted endonucleases could take advantage of the high prevalence of interspecies recombinant genotypes for UL29 and UL30 in HSV-2. These two essential genes have already been successfully targeted by CRISPR/Cas9 (van Diemen et al., 2016).

We recognize several limitations of our work. First of all, our analysis of recombinant genotypes at the UL29, UL30, and UL39 loci is limited by our sample set, which is not ideal for approximating the true prevalences of these genotypes in the populations covered. Our study constitutes a secondary analysis of HSV samples collected for various clinical studies and not at random from the represented populations. Sample sizes for geographic locations are also often limited. Because of this, the samples from any given location may represent a sub-population of the HSV-2 circulating at that location rather than being reflective of the population at large. Our results serve to put a prior probability on interspecies recombination prevalence in HSV and suggest that more rigorous sequencing of HSV from diverse human populations is required to more accurately measure this prevalence.

We also recognize that we may have underestimated the degree of HSV-1 x HSV-2 recombination, as even though the capture technique that we used for HSV sequencing greatly improved our ability to obtain high-quality, genome sequences, there remain portions of the HSV genome that are difficult to accurately sequence due to particularly high GC content and the presence of repeats. In general, these occur outside genic loci though some coding regions (such as UL36) contain difficult to sequence repeats. Because of these sequencing limitations, it remains challenging to confidently assess whether HSV-1 x HSV-2 recombination has an impact on these regions in either HSV genome.

In summary, our work demonstrates that the effects of interspecies recombination on the HSV-2 genome are less constrained than previously thought. We show that in addition to the 3 loci where recombinant genotypes are common (UL29, UL30, and UL39), there are a number of other loci that have been affected by interspecies recombination. Our results also demonstrate that recombination events can vary widely in size from about 50bp to multiple kilobases in length. We also show that interspecies recombination events continue to be generated in circulating HSV-2 clinical strains and that they are stable within hosts. Finally, we demonstrate that interspecies recombination can alter immune response to HSV by adding or removing T-cell epitopes. Together these findings suggest that interspecies recombination has significant impacts on HSV-2 genetic diversity and evolution with major implications for HSV vaccine and drug design.

## Methods

### Samples

All sequenced samples were collected from 1994 to 2016 as part of clinical research studies on HSV infection at the University of Washington Virology Research Clinic (for Seattle samples) or at international study sites in Cameroon, Peru, Senegal, and Uganda. Oral and genital swabs were either self-collected by participants or collected by clinicians. Swabs were collected directly from genital lesions or from mixed anogenital swabs as previously described (Tronstein et al., 2011). Written informed consent to collect swabs and demographic information was obtained from all participants. All studies were approved by the University of Washington (UW) Human Subjects Division and the local institutional review boards/ethics committees for the international sites.

All sequenced samples had HSV viral loads greater than 1,000 copies/mL due to limitations in recovering high confidence genomic sequences from samples with lower viral loads (Greninger et al., 2018). We selected samples in our dataset with respect to geographic origin and collection date. We also specifically included samples from individuals who had had both HSV-1 and HSV-2 detected in genital tract swabs (genital co-infection) as defined using our UL27 (gB)-based genotyping platform (Corey et al., 2005) and from individuals superinfected with more than one HSV-2 strain (as identified in Johnston et al., 2017). See Supplementary Note 3 for more details on sample selection.

### Sequencing and Consensus Generation

Samples selected for sequencing underwent the next-generation, direct-capture sequencing method for HSV previously described in Greninger et al., 2018. DNA was extracted directly from sample swabs. Pooled DNA from multiple samples was then sequenced following enrichment using HSV-1 and HSV-2 specific oligonucleotide capture panels.

Consensus sequences and .bam files for newly-generated sequences were created from raw sequencing reads using the computational pipeline as previously described in Greninger et al., 2018, which is publicly available at https://github.com/proychou/HSV/. For HSV-1 sequences, strain 17 (NC_001806) was used as the reference (Davison, 2011). For HSV-2 sequences, SD90e (KF781518) was used as the reference (Colgrove et al., 2014). Of the 70 HSV-1 and 238 HSV-2 samples we attempted to sequence, we were successfully able to generate 59 HSV-1 and 196 HSV-2 high confidence consensus sequences. Sequencing data and GenBank accession numbers for all newly-generated sequences are listed in Supplementary Tables 1 and 4. Samples we were unable to sequence are marked as “failed”.

For HSV-2, in addition to the 196 newly-generated sequences, we downloaded an additional 34 consensus sequences that had not previously been analyzed for recombination directly from GenBank. These included KY922720 – KY922726 (direct submission) and 28 sequences from Johnston et al., 2017 (see Supplementary Table 4).

### Detection of Novel Recombination Events

Each HSV-1 sequence in our dataset of 59 newly-generated sequences was aligned with HSV-2 SD90e (KF781518) and a chimpanzee herpesvirus (ChHV, NC_023677) (Severini et al., 2013) reference sequence using MAFFT (Katoh and Standley, 2013). Each of these sequence trios were then run through RDP4, version Beta 4.95 (Martin et al., 2015). The RDP program was run from the command line with the default settings. This program uses the RDP, GENECONV, Chimaera, and MaxChi algorithms to both detect recombination events and verify events identified by other algorithms (Martin and Rybicki, 2000; Padidam et al., 1999; Posada and Crandall, 2001; Smith, 1992). The algorithms BootScan, SiScan, and 3Seq are computationally intensive when used to detect new events and so are only used to verify other events when using the default settings (Boni et al., 2007; Gibbs et al., 2000; Martin et al., 2005). Putative events detected by RDP were subjected to manual review with apparently false positive events (those due to sequence misalignment or in regions of poor sequence quality) excluded from further analyses.

This process was then performed for 230 HSV-2 sequences that had not previously been analyzed for recombination with each sequence aligned to HSV-1 KOS (JQ673480) and the ChHV reference (NC_023677).

### Novel Recombination Event Verification

We confirmed novel recombination events observed in just one sample by performing Sanger sequencing across the entire event or across each of the two breakpoints, depending on the size of the event, directly from the original sample swab. Additional samples from persons with novel recombination events were sequenced when such samples were available to further confirm the recombination event and to examine the stability of the novel recombinant *in vivo*.

### Phylogenetic Trees

All phylogenetic trees were constructed using concatenated coding sequences from the genomic region of interest. Non-coding sequence was excluded due to the increased frequency of ambiguous base calls in these regions. The phylogenetic trees in Figures 1C, 1E, and 2C were created using MrBayes (Huelsenbeck and Ronquist, 2001) with a HKY85 substitution model, gamma distribution for mutation rate variation across sites with 4 rate categories, chain length of 1,100,000, and burn-in length of 100,000. PHYML with a HKY85 substitution model was used to generate the trees in Figures 1D and 2D (Guindon et al., 2010). PHYML was used in place of MrBayes for the trees in 1D and 2D given the prolonged run time required by MrBayes for alignments of long sequences.

To better illustrate the full range of genomic variation in HSV-1 and HSV-2, we used all HSV genomes available in GenBank as of April 2018 in the generation of phylogenetic trees in addition to the newly-generated genome sequences. The accessions for the GenBank sequences are listed in Supplementary Tables 1 and 4. References for these sequences are as follows: HSV-1 Colgrove et al., 2016; Davison, 2011; Greninger et al., 2018; Kolb et al., 2011; Macdonald et al., 2012a, 2012b; Parsons et al., 2015; Szpara et al., 2014; Watson et al., 2012; HSV-2 Burrel et al., 2017; Colgrove et al., 2014; Johnston et al., 2017; Koelle et al., 2017; Kolb et al., 2015; Newman et al., 2015. Within the genomic regions of interest for each tree, only one sequence per unique genotype was used. For trees constructed from alignments of relatively short sequences (1C, 1E, and 2C), sequences with any ambiguous bases were excluded. For trees constructed from alignments of longer sequences (1D and 2D), sequences with greater than 10% ambiguous base calls were excluded.

### Assignment of Genotypes for Recombinant Regions in UL29, UL30, UL39

We assigned genotypes for the UL29, UL30, and UL39 recombinant regions (as described in Burrel et al., 2017; Koelle et al., 2017) to all newly-generated HSV-2 sequences plus all those available in GenBank as of April 2018. For UL29 and UL30, we assigned genotypes based on each sample’s nucleotide sequence in the recombinant region. For UL39, genotypes were assigned based on amino acid sequence given the large number of different nucleotide genotypes observed in the region. See Supplementary Note 2 for details on how genotypes were assigned.

For the purpose of reporting the frequencies of the genotypes at the UL29, UL30, and UL39 loci, we first excluded all HSV-2 sequences with missing data at all three loci of interest. Next, we separated out samples from Burrel et al., 2017 as these samples were selected for sequencing from a larger sample set because of their unusual (non-recombinant) UL30 sequence (and therefore, the frequency of recombinant genotypes particularly at the UL30 locus in this set cannot be considered to approximate their true frequency in the represented geographic areas). We scanned these sequences for rare recombinant genotypes but did not include them in our calculations of the frequencies of common recombinant and non-recombinant genotypes. Finally, from the remaining sequences, we selected a subset which included just one sample per person. This left us with a total of 213 sequences for which we report genotype frequencies. Note that for individuals with HSV-2 superinfection (as noted in Supplementary Table 4) this means that only one of their HSV-2 strains was considered in the reported genotype frequencies.

### Structural Analyses

Depictions of the protein products of HSV-1 UL29 (ssDNA binding protein, ICP8) and HSV-1 UL30 (DNA polymerase) as determined by x-ray crystallography (Liu et al., 2006; Mapelli et al., 2005) were uploaded into Chimera (Pettersen et al., 2004), which was used to highlight the structures corresponding to genic regions affected by interspecies recombination.

We used HHPred to search for the protein with known structure that had the greatest predicted structural homology to the UL39 protein product (ribonuclease reductase) in the Research Collaboratory for Structural Bioinformatics protein database (RCSB PDB) (Berman et al., 2000). The best hit, regardless of which HSV protein sequence was used for UL39, was the large subunit of the yeast ribonuclease reductase (Xu et al., 2006). The region of this protein corresponding to the recombinant genic region was highlighted using Chimera as above.

### T-Cell Analysis

Informed written consent was obtained from subjects contributing T-cells and viral isolates. The consent process for autopsy-derived T-cells has been documented (van Velzen et al., 2013). Lab HSV-2 strains 186 (JX112656) (Anderson et al., 1980) and HG52 (JN561323), (Harland and Brown, 1985), lab HSV-1 strain E115 (Spruance and Chow, 1980), and HSV-2 sample 1996_26333 (MH790638) were grown and titered on Vero cells. Both strain 186 and HG52 have the common recombinant or CR genotype at UL30. Full-length UL30 of HSV-1 strain 17 and HSV-2 strain 186 were cloned into pDEST103 and expressed by transient transfection of Cos-7 cells as described (Jing et al., 2012; Johnston et al., 2014). For HSV-2 UL30 NR, we similarly cloned full-length UL30 from HSV-2 sample 2008_15116 (MF621257) (Johnston et al., 2017) with the same PCR primers used for HSV-2 strain 186 (Johnston et al., 2014). Accurate amplification was confirmed by sequencing and UL30 was subcloned into pDEST103 after initial cloning into pENTR221 (Jing et al., 2012). UL30 of HSV-1 17, HSV-2 186, and HSV-2 2008_15116 were also cloned into pDEST203 and expressed by in vitro transcription/translation (IVTT) as described (Jing et al., 2012). HSV-specific T-cell clones 9447.28, 1874.1991.22, and 5101.1999.23 (Supplementary Table 5) from Seattle donors have been described (Dong et al., 2010; Koelle et al., 2001, 2000). Polyclonal, mono-specific, HSV-specific CD8 T-cells were enriched from T-cells expanded from autopsy HSV-1-infected trigeminal ganglia TG3 using tetramers of HLA A*0101 and either HSV-1 UL48 AA 90-99 or HSV-1 UL48 AA 479-488, as published (van Velzen et al., 2013). Live cells positive for CD3, CD8, and tetramer were sorted (FacsAria II, Becton Dickinson) and expanded polyclonally (Koelle et al., 2001). To obtain bulk HSV-specific CD4 T-cells, HSV-1-reactive CD4 T-cells were enriched from PBMC from a Seattle HSV-1 seropositive, HSV-2 seronegative donor with no clinical history of herpes infection, as published (Jing et al., 2012). Epstein-Barr virus-transformed lymphocyte continuous lines (LCL) from persons of defined HLA type were cultured as described (Tigges et al., 1992) for use as antigen-presenting cells (APC). For functional readouts of CD8 T-cells and cloned CD4 T-cells, LCL were infected overnight with HSV and then co-cultured for an additional 24 hours with T-cells (50,000 LCL and 50,000 T-cells/well in 200 μL in triplicate). Supernatants were then assayed for IFNγ by ELISA (Koelle et al., 2001). For polyclonal CD4 T-cells, a triplicate ^3^H thymidine incorporation proliferation assay used autologous, irradiated PBMC as APC and responder T-cells (100,000 each in 200 μl/well final volumes) as published (Johnston et al., 2014). Antigens were sonicated, transfected Cos-7 cells or sonicated, UV-killed HSV-infected Vero cell preparations at final dilutions of 1:100, or IVTT protein preparations at 1:1000.

## Acknowledgements

We would like to thank the clinical study participants from whom HSV samples were collected. We would also like to thank local study teams at the international sites in Cameroon, Peru, Senegal, and Uganda. At the Senegal site, we thank Marie-Pierre Sy for swab collection. At UW, we would like to thank Chris McClurkan and Marliis Ott for assistance with T-cell experiments.

